# Generalized Reporter Score-based Enrichment Analysis for Omics Data

**DOI:** 10.1101/2023.10.13.562235

**Authors:** Chen Peng, Qiong Chen, Shangjin Tan, Xiaotao Shen, Chao Jiang

## Abstract

Enrichment analysis contextualizes biological features in pathways to facilitate a systematic understanding of high-dimensional data and is widely used in biomedical research. The emerging reporter score-based analysis (RSA) method shows more promising sensitivity, as it relies on *p-values* instead of raw values of features. However, RSA cannot be directly applied to multi-group experimental designs and is often misused due to the lack of a proper tool. Here, we propose the Generalized Reporter Score-based Analysis (GRSA) method for multi-group and longitudinal omics data. A comparison with other popular enrichment analysis methods demonstrated that GRSA had increased sensitivity across multiple benchmark datasets. We applied GRSA to microbiome, transcriptome, and metabolome data and discovered new biological insights in omics studies. Finally, we demonstrated the application of GRSA beyond functional enrichment using a taxonomy database. We implemented GRSA in an R package, ReporterScore, integrating with a powerful visualization module and updatable pathway databases (https://github.com/Asa12138/ReporterScore). We believe the ReporterScore package will be a valuable asset for broad biomedical research fields.

## Introduction

Functional enrichment analysis is a popular bioinformatic method that helps understand the biological significance of large omics datasets, such as transcriptomic, metagenomic, and metabolic data. We gain insights into the underlying biological processes and functions by identifying enriched functional categories, such as gene ontology terms or biological pathways, and formulate hypotheses for downstream experimental investigations^1^.

Methods for functional enrichment analysis can be roughly divided into three categories based on underlying statistical methods: (i) overrepresentation analysis (ORA), (ii) functional class scoring (FCS), and (iii) pathway topology-based (PT)^2^. Common enrichment analysis methods in omics research are shown in Table 1. The algorithm of reporter score-based analysis (RSA) was originally developed by Patil and Nielsen in 2005 to identify metabolites associated with the metabolic network’s regulatory hotspots^3^. The RSA has recently gained popularity due to its extended application in functional enrichment analysis in microbiome research^4^. RSA is an FCS method based on parsing the *p-values* of selected statistical analyses without a priori cut-off (threshold-free). The rationale is that the *p-value* can be considered a standardized statistic reflecting the differences between different genes or features, regardless of the mean expression values. The pathways with significantly lower *p-values* than the background *p-value* distribution are enriched^3^.

**Table 1.** Some commonly used enrichment analysis methods.

| Category | Method | Tools | Notes | Reference |
| --- | --- | --- | --- | --- |
| ORA | Hypergeometric test / Fisher's exact test | DAVID (website), clusterProfiler (R package) | The most common methods used in enrichment analysis. | 6,7 |
| FCS | Gene set enrichment analysis (GSEA) | GSEA (website), clusterProfiler (R package) | A computational method that determines whether a set of genes shows statistically significant and concordant differences between two biological states. | 8,9 |
| FCS | Generalized reporter score-based analysis (GRSA/RSA) | **ReporterScore (R package developed in this study)** | Find significant metabolites (first report), pathways, and taxonomy based on the *p-values* for multi-omics data. | 3,5 |
| PT | Reporter feature analysis | / | Integrates bio-molecular network topology with transcriptome data to identify the key biological features. | 10 |
| PT | Topology-based pathway enrichment analysis (TPEA) | TPEA (R package) | Integrates topological properties and global upstream/downstream positions of genes in pathways. | 11 |

However, RSA is often misused due to a lack of specific tools and a systematic understanding of the algorithm^5^. In addition, the sign (plus or minus) of the reporter score of each pathway in classic RSA does not represent the increasing or decreasing trend of the pathway expression; rather, reporter scores (including negative value) less than a specified threshold indicate that the corresponding pathway is not significantly enriched. This often leads to misinterpretations of the results.

Inspired by the classic RSA, we developed the improved Generalized Reporter Score-based Analysis (GRSA) method, implemented in the R package ReporterScore, along with comprehensive visualization methods and pathway databases. GRSA is a threshold-free method that works well with all types of biomedical features, such as genes, chemical compounds, and microbial species. GRSA works in the mixed (classic RSA) and directed modes (enhanced RSA). The directed mode uses signs of the reporter score to distinguish up-regulated or down-regulated pathways, which is more intuitive. Importantly, the GRSA supports multi-group and longitudinal experimental designs, because of the included multi-group-compatible statistical methods (for a full list of supported methods, please see Supplementary Table 1). The ReporterScore package also supports custom hierarchical and relational databases (e.g., table containing the correspondence between pathways and genes), providing extra flexibility for advanced users. In this study, we described the comprehensive utility of GRSA. We benchmarked GRSA against other popular enrichment methods across multiple datasets, and demonstrated the applications of GRSA on diverse omics datasets.

## Results

### Workflow overview

The ReporterScore package has built-in KEGG pathway, module, gene, compound, and GO databases and also allows customized databases, making it compatible with feature abundance tables from diverse omics data. A complete gene abundance table can be used for transcriptomic, scRNA-seq, and related gene-based omics data of a specific species. For metagenomic and metatranscriptomic data, which involve many different species, a KO abundance table can be used, generated using Blast, Diamond, or KEGG official mapper software^12^ to align the reads or contigs to the KEGG^13^ or the EggNOG database^14^. An annotated compound abundance table can be used for metabolomic data, but the standardization of compound IDs (e.g., convert compound IDs to KEGG IDs) is required.

The workflow of GRSA in the ReporterScore package is shown in Fig. 1, using metagenomic data as an example. The KO abundance table (rows are KOs and columns are samples) and metadata table (rows are samples and columns are experimental design groups) were used as the input for GRSA. Importantly, the input data should not be prefiltered to retain the background information. First, the *p-values* for all KOs were calculated by a proper statistical method. Then, in the classic mode, the *p-values* were directly converted to Z-scores (Fig. 1a, equation [i]). In the directed mode, the *p-values* were divided by 2, converted to Z-scores, and assigned plus or minus signs, denoting up- and down-regulated KOs (Fig. 1a, equation [ii-iv]). Next, the Z-score of the pathway 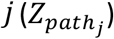 was calculated by summing the Z scores of KOs within the pathway *j* and dividing by the square root of the number of KOs (*k*_*j*_) in the pathway *j* (Fig. 1a, equation [v]). The 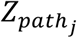 is further standardized by the background pathway Z-score distribution, generated by randomly sampling *k*_*j*_ KOs from the total KO pool (Fig. 1a, equation [vi]). The standardized pathway Z-score is defined as the reporter score of a pathway (*ReporterScore*_*j*_). The details of the GRSA algorithm are described in the Methods section.

**Fig. 1.**
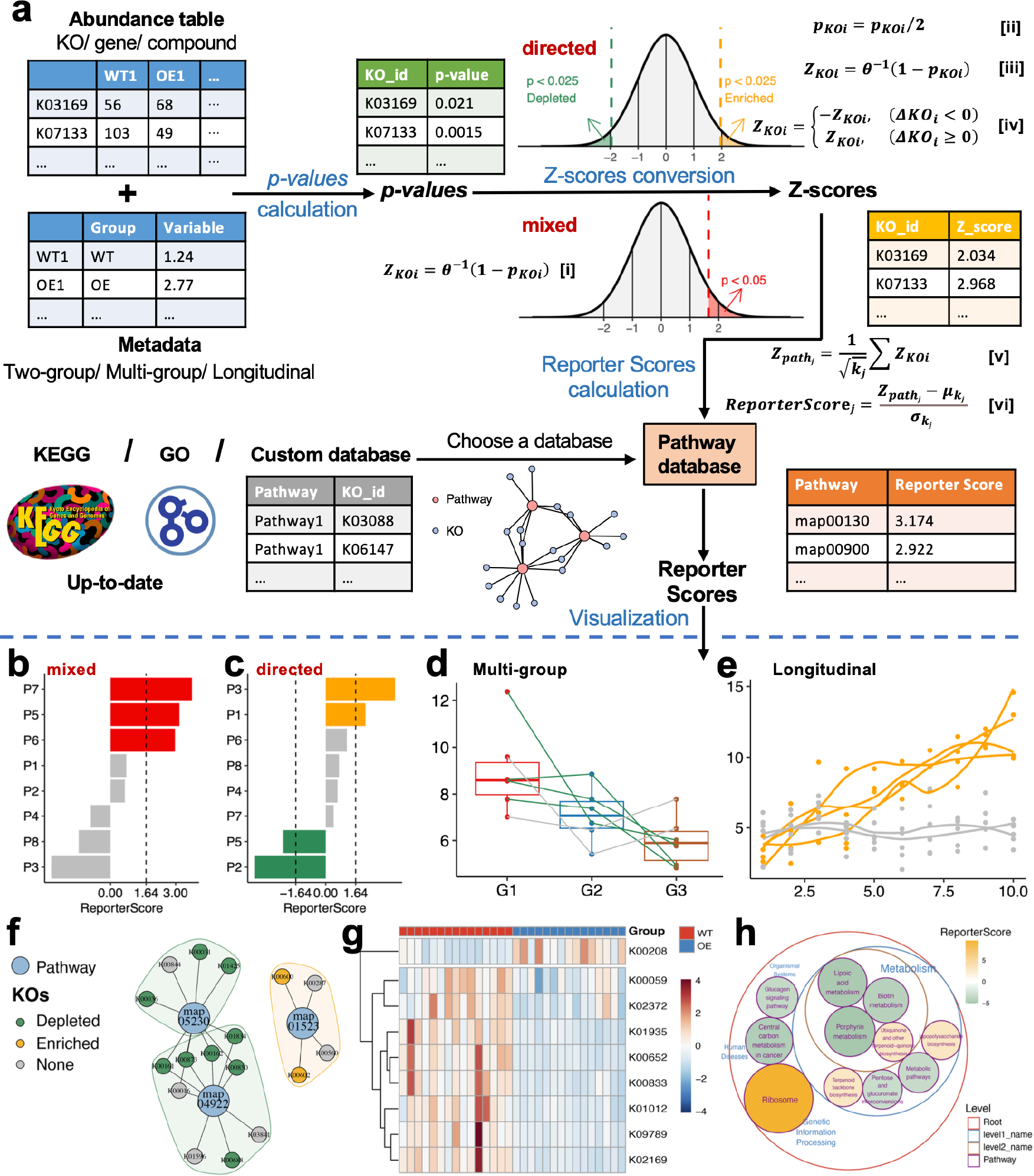
The overall workflow of GRSA in the ReporterScore package. **a** GRSA workflow consists of four parts: Calculate the *p-values* for KOs between two or multiple groups by various statistical methods; convert the *p-values* of KOs to Z-scores by inverse normal distribution and assignment of a plus or minus sign to each Z-score in the directed mode; mapping KOs to annotated pathways and calculating the reporter score for each pathway; visualize the results of GRSA. *KO*_*i*_ represents a certain 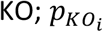 is the *p-value* of *KO*_*i*_ ; 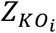 is the Z-score transformed from 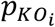; *ΔKO*_*i*_ is the abundance difference of between groups. A total of *k*_*j*_ KOs were annotated to the corresponding pathway. 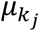 and 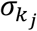 are the mean and the standard deviation of the background Z-score distribution *Z*_*path_null*_, respectively. The ReporterScore package provides various visualization methods for the GRSA result: **b** The bar chart shows reporter scores of pathways in the mixed mode. The red color indicates significantly enriched pathways, with reporter scores greater than 1.64, corresponding to a *p-value* of 0.05. **c** The bar chart shows reporter scores of pathways in the directed mode. The orange and green colors indicate up-regulated and down-regulated pathways with absolute reporter scores greater than 1.64. **d**-**e** The line charts show the pattern of a selected pathway in the directed mode with a multi-group (**d**) and a longitudinal design (**e**), each line represents the trend of the average abundance of one KO. Line colors indicate whether the KO is significantly enriched (orange), depleted (green), or neither (grey). **f** The network plot shows the KOs present in selected pathways; some KOs can be shared by several pathways. Big dots represent pathways, and small dots represent KOs. The colors of small dots represent the trend of KOs. The colors of the shades encircling pathways denote whether the pathway is overall up-regulated (orange) or down-regulated (green). **g** The heatmap displays the abundance of each KO in a pathway for different samples (columns). **h** The circular packing plot shows the hierarchical relationship of selected pathways; the size of the circle indicates the absolute value of the reporter score, and the color of the circle indicates that the pathway is overall up-regulated (orange) and down-regulated (green).

We designed the ReporterScore package to be user-friendly. In one step, the function `reporter_score` calculates the reporter scores for a feature abundance table with associated metadata. The included assorted visualization methods can be used to explore the entire pathways and features within pathways (Fig. 1b-h). The demo code is included in the Methods section.

### Applying GRSA to multi-group and longitudinal omics data

An important feature of GRSA is the newly developed directed mode. The key difference between the directed mode and the mixed mode (classic RSA) is that in the directed mode, the reporter score’s plus or minus sign indicates the pathway’s increasing or decreasing trend (Fig. 1c). However, in the mixed mode, the signs of the reporter score do not indicate the trends of the pathways. We performed GRSA on the public ex_KO_profile dataset (a metagenomic dataset) in two modes (Supplementary Fig. 1a). For pathways enriched in the directed mode, most KOs within the pathway shared the same trend. The pathway with consistently increasing (decreasing) KOs would acquire a significantly bigger (smaller) aggregated Z-score than the background (Supplementary Fig. 1b, blue and red boxes). If KOs within a pathway had opposing trends, the signed Z-scores would cancel each other, leading to insignificant results (Supplementary Fig. 1b, orange box). In comparison, in the mixed mode, the increasing and decreasing trend of the enriched pathway cannot be determined (Supplementary Fig. 1c). Therefore, the directed mode helps find pathways with consistently changing KOs. For simplicity, we used GRSA in the directed mode henceforth.

Another major advantage of GRSA is the full support of multi-group and longitudinal omics data. The ReporterScore package calculates the *p-value* for each feature between groups using differential abundance analysis or correlation analysis. The Kruskal-Wallis test or ANOVA assesses if the feature abundance varies significantly across multiple groups. The default correlation analysis treats group assignments as ordinal (e.g., groups “G1”, “G2”, and “G3” will be converted to 1, 2, and 3), so the correlation analysis could evaluate if the feature abundance linearly increases or decreases. Moreover, the ReporterScore package also supports any specified patterns. For example, groups “G1”, “G2”, and “G3” can be set as 1, 10, and 100 if an exponentially increasing trend is expected. To explore potential patterns within the data, clustering methods, such as C-means clustering, can be used.

As a general rule, the users must ensure the selected statistical methods are applicable to the datasets and experimental designs. We applied GRSA with different statistical methods on multiple benchmark datasets. For the classic two-group design, the Jaccard similarity exceeded 0.84 for parametric methods and 0.78 for non-parametric methods, but the Jaccard similarity between parametric and non-parametric methods was lower than 0.63 (Supplementary Fig. 2a). The differences mainly stemmed from parametric versus non-parametric methods. For the multi-group data, users can choose the differential abundance analyses if they aim to enrich significantly altered pathways across groups. Correlation analysis is the preferred choice if the goal is to enrich pathways that show consistent increasing or decreasing patterns (Supplementary Fig. 2b,c). Lastly, GRSA also supports other statistical tests, such as ‘DESeq2’, ‘Edger’, ‘Limma’, ‘ALDEX’, and ‘ANCOM’^15^, to calculate the reporter scores. The demo code is shown in the Methods section.

### GRSA showed higher sensitivity than other commonly used enrichment analysis methods

Next, we evaluated the performance of GRSA and compared it with other commonly used enrichment analysis methods (Table 1) on several benchmark datasets (Supplementary Tables 2 and 3). PT-based methods may better identify biologically meaningful pathways than non-PT-based methods in certain scenarios^16^. However, PT-based methods require comprehensive topological structures of pathways, limiting their applications in other non-human organisms^17^. Therefore, we focused on the comparison against non-PT enrichment analysis methods.

Nguyen et al. proposed several approaches to compare enrichment methods^16^, we adapted their methodologies and evaluated the performance of GRSA against the other popular enrichment analysis methods using the identical pathway database: Fisher’s exact test (`fisher.test`; R base package), an improved form of Fisher’s exact test (`enricher`; `clusterProfiler` package), and gene set enrichment analysis (GSEA).

We first compared the efficacy of different methods in identifying the target disease pathways using the 24 gene expression datasets associated with known human diseases (Supplementary Table 2). The rationale is that since each dataset is affiliated with a specific disease-linked KEGG pathway (the target pathway), an optimal enrichment analysis method should rank the target pathway among the top of all 342 pathways and enrich the target pathway with a smaller adjusted *p-value*. Notably, GRSA exhibited the highest performance among the four methods (Fig. 2a,b). Generally, threshold-free methods, such as GRSA and GSEA, outperformed the conventional ORA methods.

**Fig. 2.**
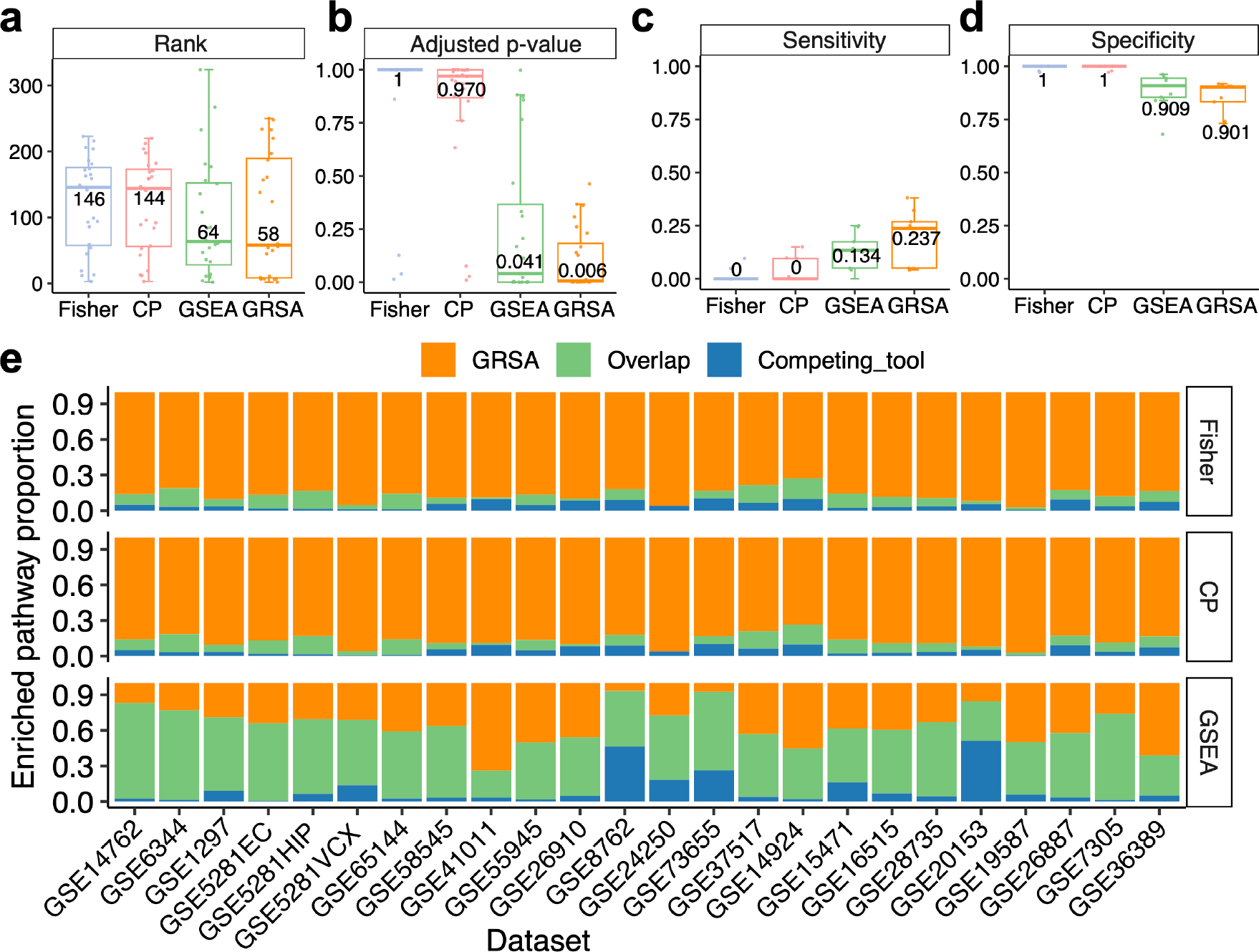
Comparisons of GRSA and other commonly used methods of enrichment analyses. **a**-**b** The box charts show the ranks (**a**) and adjusted *p-values* (**b**) of target pathways derived by four methods on 24 gene expression datasets. Numbers represent the median values for each method. **c**-**d** The box charts show the sensitivity (**c**) and specificity (**d**) of four methods on nine wild-type/knockout gene expression datasets. Numbers represent the median values for each method. **e** Proportion of enriched pathways in GRSA and other enrichment analysis methods based on 24 gene expression datasets. Orange pathways were only identified by GRSA, blue pathways were only identified by the competing tool and green pathways were identified by GRSA and the competing tool. Fisher: fisher.test; CP: improved fisher.test used by clusterProfiler; GSEA: gene set enrichment analysis by clusterProfiler; GRSA: generalized reporter score-based analysis.

We next evaluated the proficiency of different methods in detecting pathways disturbed by gene knockout experiments. In a gene knockout experiment, the knockout gene is a confirmed source of perturbation. Nguyen et al. considered pathways that included the knockout genes as true positives and pathways not including the target genes as true negatives, so the enrichment of such pathways would be considered false positives. Under these assumptions, we could calculate the sensitivity and specificity of a method. GRSA showed the highest median sensitivity among the methods considered, although its specificity is slightly reduced compared to others (Fig. 2c,d). We prioritize the sensitivity of a method because for pathways that included the knockout gene, deleting it should have a sizable impact on the pathway (sensitivity); however, for pathways that do not include the genes in the database, given the potential incompleteness of pathway and gene databases, simply attributing these enriched pathways as false positives (specificity) may not always be appropriate.

Lastly, we evaluated the ability of different methods to enrich biologically meaningful pathways^11^. We compared the proportions of pathways identified by the GRSA, the competing tool, and both, using the number of all significant pathways as the denominator (Fig. 2e). GRSA consistently identified a larger proportion of pathways than the two ORA methods and largely overlapped with GSEA. In 21 of 24 gene expression datasets associated with human diseases, GRSA identified more enriched pathways than GSEA. For example, in the renal cell carcinoma dataset (GSE6344), pathways related to cytokine−cytokine receptor interaction^18^, IL−17 signaling^19^, and PI3K−Akt signaling^20^ were only enriched by GRSA (Supplementary Fig. 3a). In the endometrial cancer dataset (GSE7305), GRSA identified pathways related to cancer, Toll−like receptor^21^, and cortisol synthesis^22^, which have been shown to be involved in the pathological characteristics of endometrial cancer (Supplementary Fig. 3b). Therefore, GRSA can potentially identify more pathways biologically relevant to the studied diseases.

### Case Study 1: The functional analysis and age-related dynamics of the skin microbiota

Next, we showcased the versatile applications of GRSA with different types of omics data. For microbiome data, we collected the KO profile of the IHSMGC (integrated Human Skin Microbial Gene Catalog) dataset published by Wang et al.^23^ and re-analyzed the data using the GRSA method. The previous study calculated the pathway abundance by aggregating the abundances of features within a pathway and then performed differential abundance analysis to identify the most significantly differentially abundant functional modules related to cutotypes. We applied the GRSA to find the functional differences between the two cutotypes. The results were largely consistent. As an example, modules related to the biosynthesis of thiamine, phylloquinone, and cobalamin were enriched in the *M-cutotype*, while modules related to tetrahydrofolate, menaquinone, pantothenate, and ubiquinone were enriched in the *C-cutotype* (Fig. 3a). In addition, the *M-cutotype* was enriched with a large number of modules related to the metabolism of sulfur, phenylacetate (aromatic compound), and amino acids, while the *C-cutotype* was enriched with modules related to carbohydrate metabolism (Supplementary Fig. 4). Importantly, GRSA also identified pathways not found in the previous study. The *M-cutotype* was enriched with modules related to nucleotide metabolism, such as the degradation and de novo biosynthesis of purine (Supplementary Fig. 4), indicating that the *M-cutotype* microbiota may have a higher nucleotide turnover rate and stronger proliferation^24^.

**Fig. 3.**
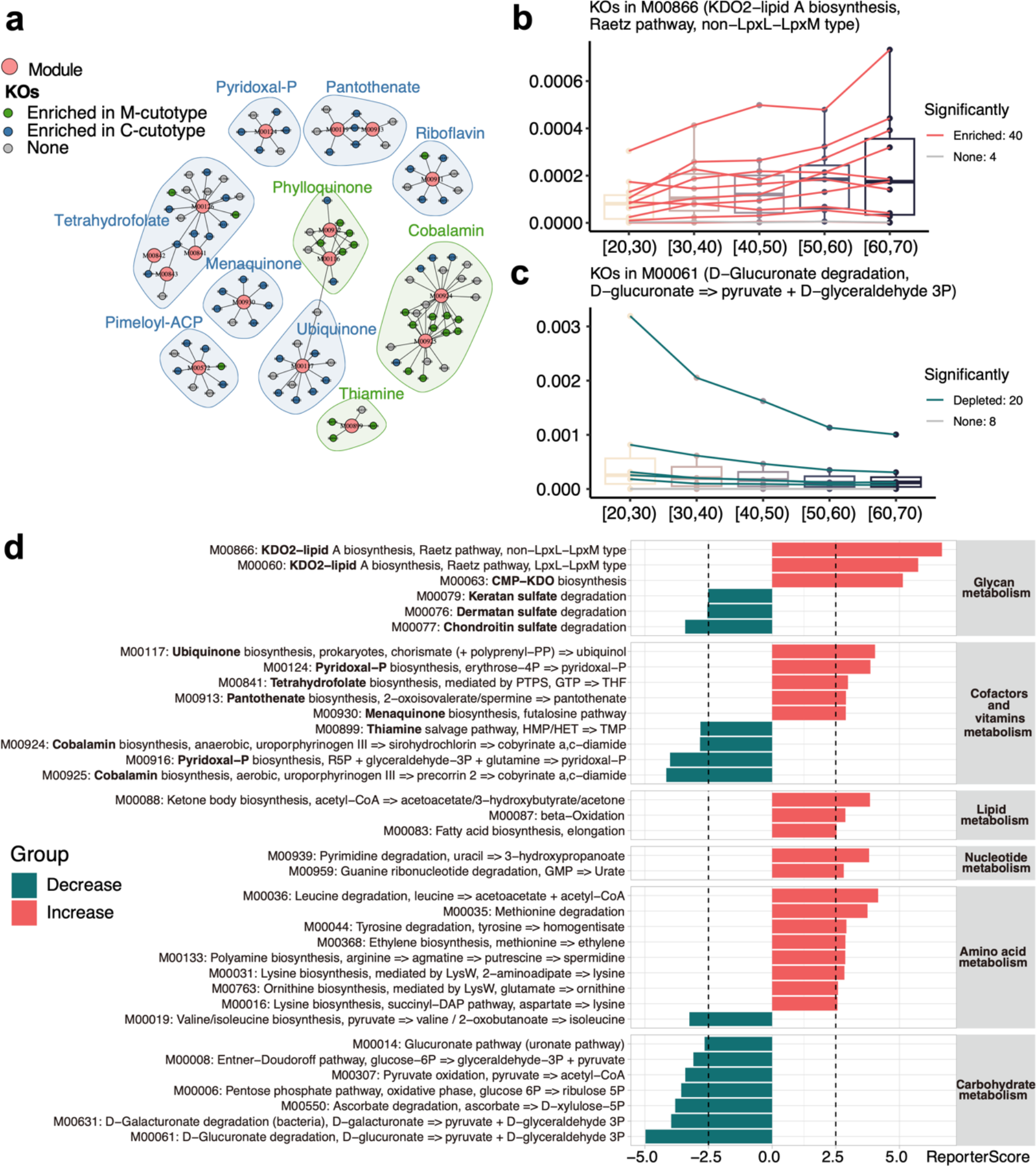
Application of GRSA to the skin microbiome of the IHSMGC dataset. **a** The network of KO-Module enriched in the *M-cutotype* (green) and *C-cutotype* (blue). Only modules related to vitamin biosynthesis were shown. Big dots represent modules; small dots represent KOs. The colors of small dots represent cutotypes. Shades indicate modules involved in the biosynthesis of the same vitamin. The colors of shades denote modules enriched in the *M-cutotype* (green) or enriched in the *C-cutotype* (blue). **b**-**c** The Box charts of modules ‘M00866’ (**b**) and ‘M00061’ (**c**) across ages. The colors of the lines represent the trend of KO’s relative abundance in the module. ‘M00866’ had the biggest positive reporter score (increasing), while ‘M00061’ had the biggest absolute value of negative score (decreasing). **d** The Bar charts show significantly enriched modules over aging; the reporter score threshold of 2.5 corresponds to a confidence of about 0.995, and these modules are grouped based on the KEGG level B. Colors denote up-regulated (red) or down-regulated (green) modules with aging.

The previous study divided samples into five aging groups and found that the prevalence of the *M-cutotype* significantly increased with age. However, they did not perform age-related functional analysis. We re-analyzed the multi-group data using GRSA to explore the functional dynamics related to aging. The larger positive reporter scores indicate that the module had an overall increasing trend with respect to age, such as ‘M00866’, related to lipid A biosynthesis (Fig. 3b), while modules with negative reporter scores show an overall decreasing trend with respect to age, such as ‘M00061’, related to D-Glucuronate degradation (Fig. 3c). We next analyzed the chronological trend of the functional modules at the KEGG level B (Fig. 3d), which better reflects the overall metabolic activities of the microbiome. We found that the carbohydrate metabolism activity of the skin microbiota decreased with aging, while the lipid, amino acid, and nucleotide metabolism activity increased with aging. These results suggest that the energy sources of the skin microbiota significantly change with aging.

The vitamin biosynthesis-related functional modules also showed differences with respect to aging (Fig. 3d). For the glycan metabolism-related functional modules, biosynthesis of KDO2-lipid A and CMP-KDO increased with aging. KDO2-lipid A is an essential lipopolysaccharide (LPS) component in most gram-negative bacteria, which has endotoxin activity and stimulates host immune responses through Toll-like receptor 4 (TLR4)^25^. CMP-KDO is an important intermediate in the synthesis of KDO2-lipid A, and CMP-KDO synthesis is the key rate-limiting step for introducing KDO into LPS^26^. These results suggest that the microbiota of aging skins likely accumulated endotoxins, which can stimulate host inflammation. In addition, we found degradation pathways of several sulfated glycosaminoglycans (chondroitin sulfate, dermatan sulfate, and keratan sulfate) decreased in aging skin. Sulfated glycosaminoglycans play a key role in regulating skin physiology, and there is ample evidence that their properties and functions change over time and with extrinsic skin aging^27,28^. Total sulfated glycosaminoglycan abundance was reduced in aging skin^29^, which may lead to the decreased degrading ability of the skin microbiota for the sulfated glycosaminoglycans.

### Case Study 2: The functional transcriptional dynamics during cardiomyocyte differentiation

We applied GRSA to the transcriptomic dataset published by Liu et al. in 2017^30^. The study used the WGCNA method to analyze the temporal transcriptomic changes during the differentiation of cardiomyocytes from 2 hiPSC lines and 2 hESC lines at 4 timepoints (pluripotent stem cells at day 0, mesoderm at day 2, cardiac mesoderm at day 4, and differentiated cardiomyocytes at day 30). Significant changes were observed in the four stages of differentiation among all cell lines. For example, genes in module 1 were highly expressed only in differentiated cardiomyocytes (stage CM), and their enriched Gene Ontology (GO) terms of Biological Process (BP) were related to heart functions, such as regulation of cardiac contraction and muscular system processes. However, WGCNA did not assume the patterns to be linear so that genes can be only highly expressed at day 2 during mesoderm development, for example.

In addition to linearly increasing or decreasing patterns, GRSA allows users to specify any expected patterns for enrichment analysis. To start, we used the fuzzy C-means clustering method to identify the main gene expression patterns (Fig. 4a) and then used these patterns for GRSA to obtain significantly enriched pathways in each pattern (using the RSA_by_cm function in the ReporterScore package). For example, ‘Heart process (GO:0003015)’ was a significantly enriched GO term for Cluster 6, which was highly expressed only in stage CM (day 30). We identified many genes consistent with the expression pattern of Cluster 6 (Fig. 4b).

**Fig. 4.**
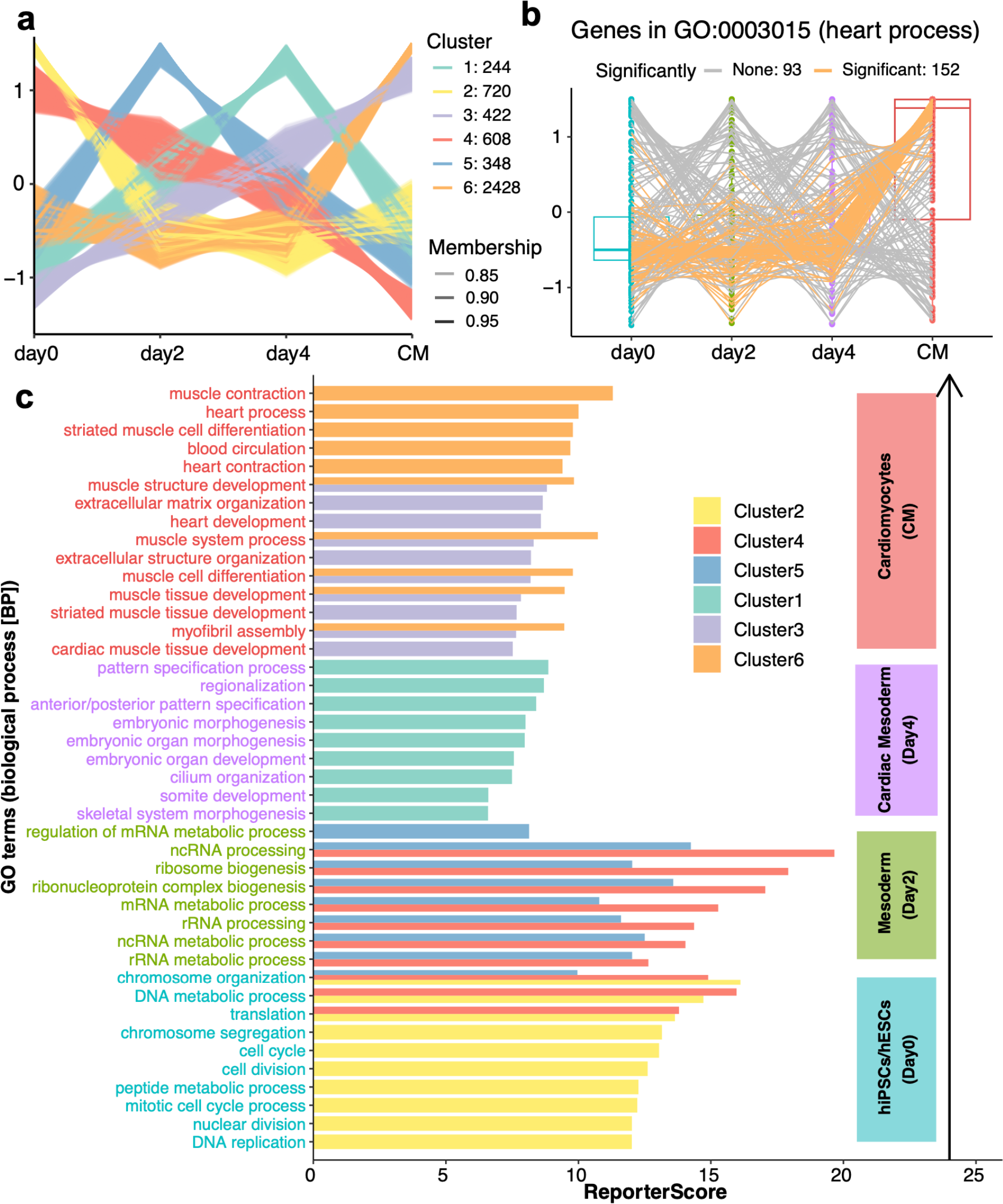
Application of GRSA to the transcriptomic dataset on the cardiomyocyte differentiation processes. **a** C-means clustering result of gene abundance profiles across four differentiation stages. The genes with membership scores greater than 0.8 were displayed. The alpha (transparency) of each line was related to the value of its membership score, and the y-axis represent the standardized abundance. **b** The box chart shows the abundance of genes in ‘GO:0003015’ (heart process) across four time points; the colors of the lines represent the correlative significance of each gene with Cluster 6 within the GO term. ‘GO:0003015’ is a representative term of Cluster 6. **c** The bar chart shows significantly enriched GO terms for each clustering pattern corresponding to the differentiation stages. The colors of the bars represent the cluster information, and the representative GO terms with high reporter scores in each cluster were shown. The text labels on the left were colored according to the stages with the highest expression. In general, Cluster 2 corresponds to day 0, Clusters 4 and 5 correspond to day 2, Cluster 1 corresponds to day 4, and Clusters 3 and 6 correspond to CM. Note that only pathways with significant positive scores are shown. The negative score of specified patterns would indicate anti-correlative patterns, which should have already been identified by c-means analysis, such as cluster 3 vs. 4.

GRSA results for all clusters are shown in Fig. 4c. Cluster 2 was highly expressed only at day 0, and its enriched GO terms were mainly related to the mitotic cell cycle, which was expected for stem cell self-renewal processes. Cluster 5 had the highest expression level on day 2 and was mainly enriched in various transcription and translation processes. Many known transcription factors, such as EOMES, MIXL1, and WNT3A, were assigned to this cluster and play important roles in mesoderm development. Cluster 4 was highly expressed at day 0 and day 2 and showed a gradually decreasing trend, its function overlapped with Clusters 2 and 5. Cluster 1, highly expressed at day 4, was related to mesoderm formation, such as morphogenesis and organ development. Clusters 3 and 6 were primarily up-regulated in differentiated cardiomyocytes (CM stage), and they were related to heart functions, such as regulation of heart contraction and muscle system processes, similar to module 1 in the previous study. Interestingly, the biological processes of hiPSCs/hESCs at day 2 (Cluster 5) induced various RNA-related metabolisms, which were not found in the previous study, indicating that complex transcriptional regulations are involved for further mesoderm formation. Therefore, using the identified expression patterns across groups, we successfully identified pathways and modules essential to different stages of the cardiomyocyte differentiation processes.

### Case Study 3: The systematic maternal metabolomic changes correlated with gestational age

We next applied GRSA to metabolomic data from a Danish pregnancy cohort in which female participants had blood drawn weekly from pregnancy to the postpartum period for untargeted metabolomics analysis^31^. Using gestational age as the study variable, they modeled a metabolic clock and found that several marker metabolites increased linearly with gestational age.

We found several important pathways upregulated with gestational age: steroid hormone biosynthesis, cortisol synthesis and secretion, and Oocyte meiosis (Fig. 5a). Multiple steroid hormones were up-regulated with increasing gestational age (Fig. 5b), including progesterone that interacts with the hypothalamic-pituitary-adrenal axis (HPA axis)^32^ and estriol-16-glucuronide produced by the placenta^33^. At the same time, two androgen-related steroid hormones were down-regulated: dehydroepiandrosterone sulfate and androsterone 3-glucuronide, as the concentration of androgens plays important physiological functions during pregnancy^34^. We also found that pathways related to the metabolism of aromatic amino acids were down-regulated with increasing gestational age (Fig. 5a), which has been reported^35^.

**Fig. 5.**
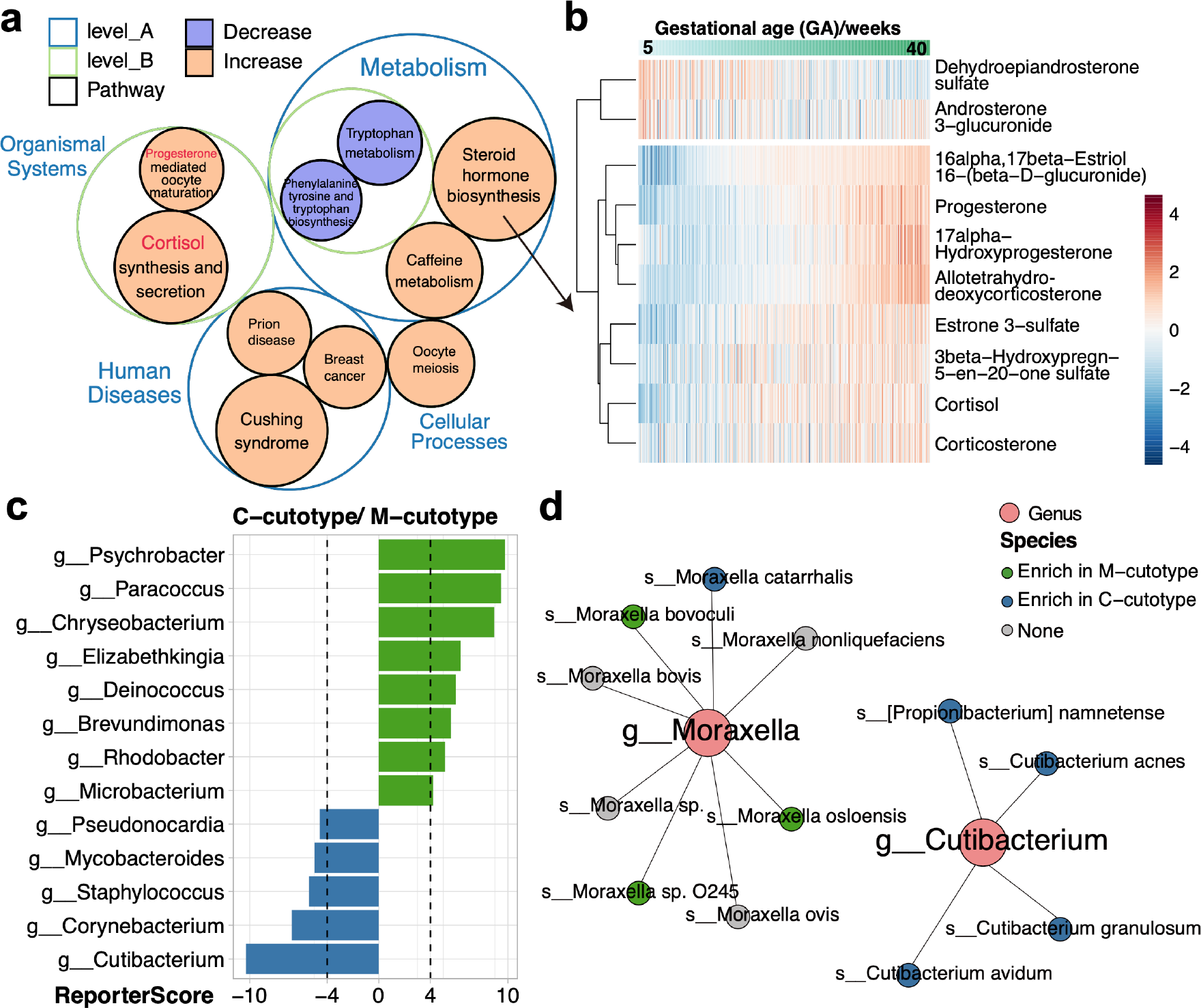
Application of GRSA in the metabolic data of the Danish pregnancy cohort and the taxonomic enrichment analysis of the IHSMGC dataset. **a** The circle packing chart shows the hierarchical relationships of significantly enriched pathways identified by GRSA in the metabolomic study. The size of the circle indicates the absolute value of the reporter score, and the color of the circle indicates the sign of the reporter score. The positive reporter score indicates the pathway was increased (orange), while the negative reporter score indicates the pathway was decreased (purple). **b** The heatmap shows the abundance of metabolites in the pathway ‘steroid hormone biosynthesis’. The columns are samples ordered by the increasing gestational age. **c** The bar chart shows significantly enriched genera in the *C-cutotype* and *M-cutotype*. **d** The network plot shows the species in *g_Moraxella* and *g_Cutibacterium* enriched in the *M-cutotype* (green) or *C-cutotype* (blue).

Importantly, we identified several up-regulated pathways related to human diseases not mentioned in the previous study. Cushing syndrome happens when the body has too much of the hormone cortisol for a long time, which could be induced by a healthy pregnancy^36^. The up-regulation of pathways related to breast cancer was also noticeable, as Pregnancy-Associated Breast Cancer (PABC) accounts for 7% of all breast cancer in young women^37^. More potential discoveries can be made if the metabolite-pathway database is improved.

### Case Study 4: The application of GRSA beyond functional enrichment analysis

The algorithm of GRSA suggests that any features organized in a hierarchical relationship can be used as an enrichment database. For example, we can perform taxonomic enrichment analysis using the phylogenetic relationships of microbes, such as genus-species relationships. To demonstrate, with the `custom_modulelist` function, we used the species abundance table from the IHSMGC dataset and looked for genera enriched in the two cutotypes. We found that *Psychrobacter, Paracoccus, Chryseobacterium, Elizabethkingia, Deinococcus*, and *Microbacterium* were enriched in the *M-cutotype*, while *Acidipropionibacterium, Staphylococcus, Corynebacterium*, and *Cutibacterium* were enriched in the *C-cutotype* (Fig. 5c), some of which were highly consistent with the differential species modules found by co-occurrence network in the previous study. However, we additionally found some genera, such as *Brevundimonas* and *Rhodobacter*, enriched in the *M-cutotype*, while *Pahexavirus* (phages of *Propionibacterium* and *Cutibacterium*) was enriched in the *C-cutotype* (Fig. 5c), probably due to the higher sensitivity of GRSA.

Two species, *Moraxella osloensis* and *Cutibacterium acnes*, were used to define the cutotype in the previous study. Interestingly, while the *Cutibacterium* genus was a good biomarker between cutotypes, the *Moraxella* genus was not, as the included species did not share the same trend (Fig. 5d). Therefore, in addition to functional enrichment analysis, GRSA can be extended to any hierarchical relational data structure.

## Discussion

We developed the ReporterScore package and demonstrated broad applications of the GRSA enrichment analyses. We improved the classic RSA method for easier interpretation of the plus and minus signs of the reporter scores. More importantly, we expanded the scope of GRSA from two-group designs to multi-group and longitudinal designs. We demonstrated these new features with metagenomic, transcriptomic, and metabolomic data (Fig. 3-5). Lastly, we showed that the GRSA is not limited to functional enrichment analysis. Notably, all figures were generated using the visualization module in the ReporterScore package.

GRSA considers all KOs involved in the pathway compared to hypergeometric tests that only consider a pre-defined list (e.g., KO/gene with *p-value* < 0.05). Thus, GRSA is more sensitive and can comprehensively assess the feature abundance differences in the pathway (Fig. 2). Compared with GSEA, which considers the magnitude of gene changes for ranking, GRSA uses *p-values* for ranking and permutation. We considered sensitivity a crucial metric for enrichment methods because an effective method should maximize the identification of the expected pathways (Fig. 2c). In contrast, we cannot precisely define the true negative results due to the incompleteness of databases. Thus, the specificity of a method cannot be accurately determined (Fig. 2d). Moreover, the primary role of functional enrichment analysis is to guide for interpreting the omics data. Additional experimental verification is essential to substantiate the analysis results.

Multi-omic studies are increasingly prevalent and so are the demands for functional enrichment analysis of all types of omics data^38,39,40,41^. The GRSA applies to all types of omics datasets as long as a relevant relational database is available. As demonstrated by case studies, we confirmed previous key findings and acquired new biological insights. For example, applying GRSA on the IHSMGC dataset suggested different functional profiles between aging and young skin microbiota. Biosynthesis of KDO2-lipid A and CMP-KDO increased while degradation pathways of several sulfated glycosaminoglycans decreased in older skin microbiota, which may be linked to changes in the skin’s physiological properties. Further studies are needed to investigate the underlying mechanisms and their implications for skin health. Applying GRSA on the transcriptomic data of cardiomyocyte differentiations revealed that hiPSCs/hESCs at day 2 specialized in various RNA-related metabolisms, suggesting the involvement of complex transcriptional regulation in further mesoderm formation. Finally, applying GRSA to the metabolomic data from the Danish pregnancy cohort showed that several pathways related to human diseases were up-regulated with gestational age, including Cushing syndrome and PABC.

The GRSA offers the option for user-specified patterns for enrichment analysis, allowing for rapid testing of educated hypotheses in complex multi-group studies. This is demonstrated in our analysis of the transcriptomic data during cardiomyocyte differentiations. The GRSA offers applications beyond functional enrichment analysis. Applying GRSA in the taxonomic enrichment analysis of the IHSMGC dataset identified key genera that significantly differed between the two cutotypes. The results were highly consistent with the microbial co-occurrence network analysis in the previous study, but performing GRSA was much easier and faster than the network analysis.

In summary, we believe the GRSA and the ReporterScore package can greatly facilitate the functional enrichment analyses of diverse omics data, with higher sensitivity, compatibility with multi-group and longitudinal designs, and flexibility with customized databases for creative applications beyond functional enrichment analyses.

## Methods

### Algorithm

The algorithm of GRSA is described as follows, using metagenomic data as an example.

#### (1) Calculating the *p-values*

A statistical method (the full list of supported statistical methods is in Supplementary Table 1) was used to obtain the *p-values* of the features (i.e., 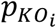, *KO*_*i*_ represents a certain KO; Fig. 1a). We used KO to represent different features in the formulas for simplicity.

#### (2) Converting the *p-values* into Z-scores

For the classic mixed RSA, we used an inverse normal cumulative distribution function (*θ*^−1^) to convert the *p-values* of KOs into Z-scores. Thus, assuming uniformly distributed *p-values* under the random data assumption ranges, the resulting Z-scores will follow a standard normal distribution (Fig. 1a), the formula is:

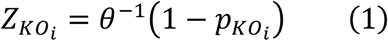

For the new directed RSA, we first divided the *p-values* by 2, transforming the range of *p-values* from (0,1] to (0,0.5]:

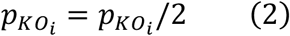

Secondly, we used an inverse normal cumulative distribution function to convert the *p-values* of KOs into Z-scores (equation (1)). When the *p-value* is 0.5, the converted Z-score equals 0. Since the above *p-values* are no greater than 0.5, all converted Z-scores will be greater than 0 (Fig. 1a).

We then determined if a KO is up-regulated or down-regulated and calculated the *ΔKO*_*i*_. In a differential abundance analysis (two-group design):

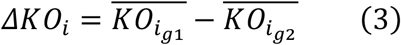

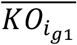 is the average abundance of *KO*_*i*_ in group1, and 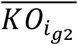 is the average abundance of *KO*_*i*_ in group2.

In a correlation analysis (two-group, multi-group, and longitudinal design):

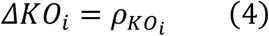

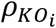 is the correlation coefficient between *KO*_*i*_ and the numeric variable. Finally, assign a plus or minus sign to each Z-score:

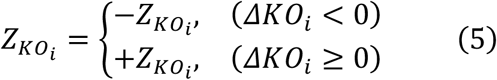

Therefore, a *KO*_*i*_ with a 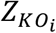 greater than 0 is up-regulated, a *KO*_*i*_ with a 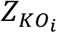 less than 0 is down-regulated.

#### (3) Scoring the pathway

We next used the Z-scores of KOs to score the pathway. First, choose a pathway database as the reference. It is of particular interest to note any hierarchy relational table (e.g., KEGG, taxonomy database) can be used as a reference as long as the relationship between the upstream and downstream features (e.g., pathways and KOs) can be represented by a bipartite network (Fig. 1a). For each pathway in the selected database, calculate the Z-score of pathway 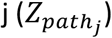 as follows:

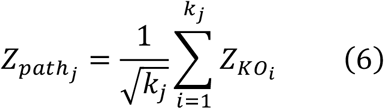

where 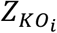 is the Z-score of *KO*_*i*_ within the *path*_*j*_, and *k*_*j*_ denotes the total number of KOs in the *path*_*j*_;

Next, we corrected 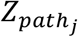 using the randomly sampled 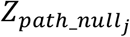 from the background distribution of the Z-scores of all 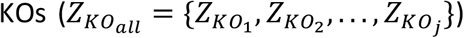 to evaluate the significance of enrichment. Specifically, for a given pathway *path*_*j*_ including *k*_*j*_ KOs, we randomly sampled the same number (*j*) of KOs from the background 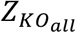 without replacement, and calculated the 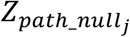 for N times (N = 10,000 in this study^10^). We then standardized 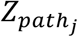 by subtracting the mean 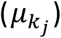 and dividing by the standard deviation 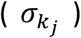 of the 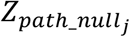 distribution. The standardized 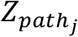 is the *ReporterScore*_*j*_. The *p-value* of *ReporterScore*_*j*_ is also estimated by the above permutation. The formulas for the reporter scores and associated *p-values* are:

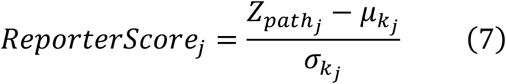

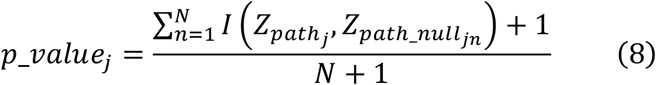

where 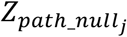 have the same *k* to 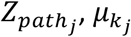 is the mean of the randomly generated N 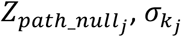 is the standard deviation of the randomly generated N 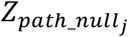.

For the classic mixed RSA:

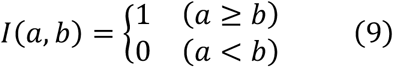

For the new directed RSA:

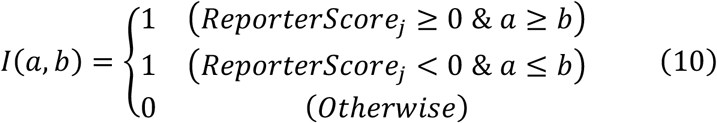

### Demo codes

An example code tailored for one step GRSA of a KO abundance table is as follows.

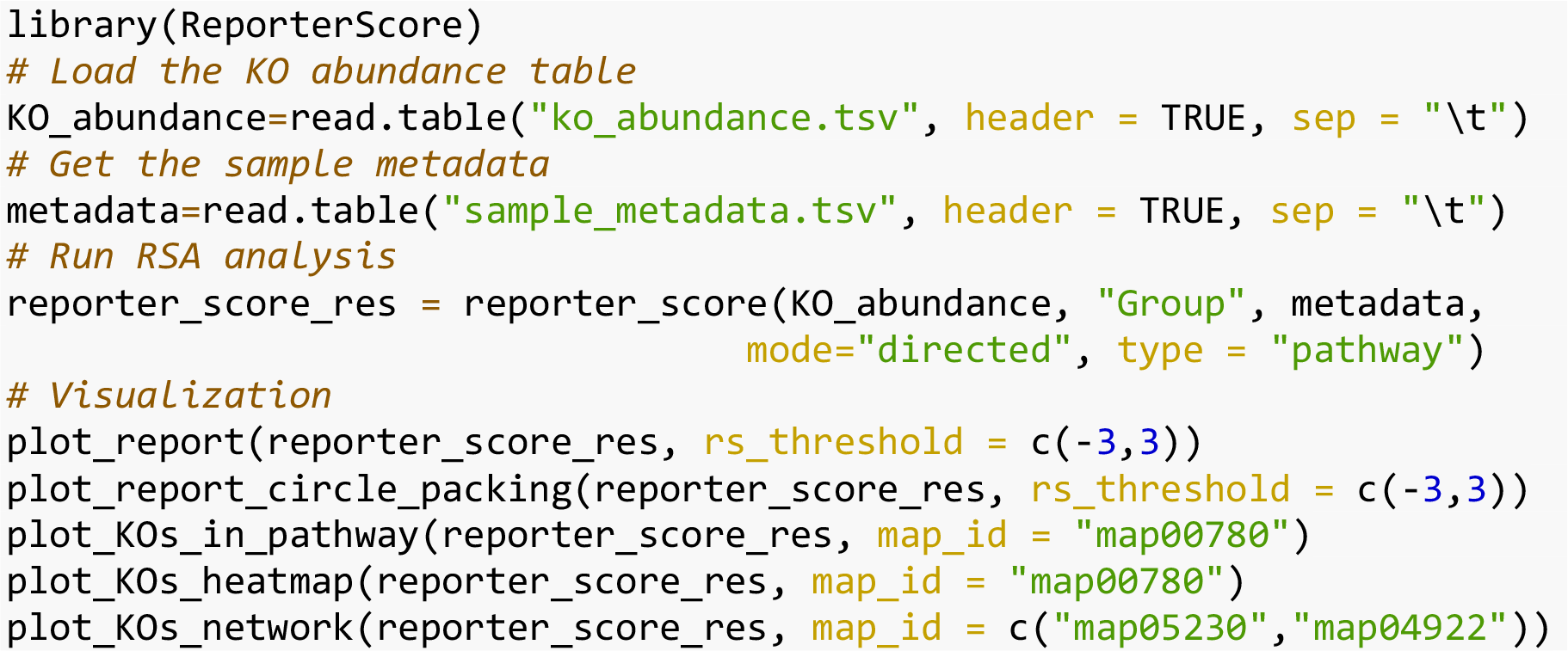

An example code for GRSA using other statistical tests is as follows.

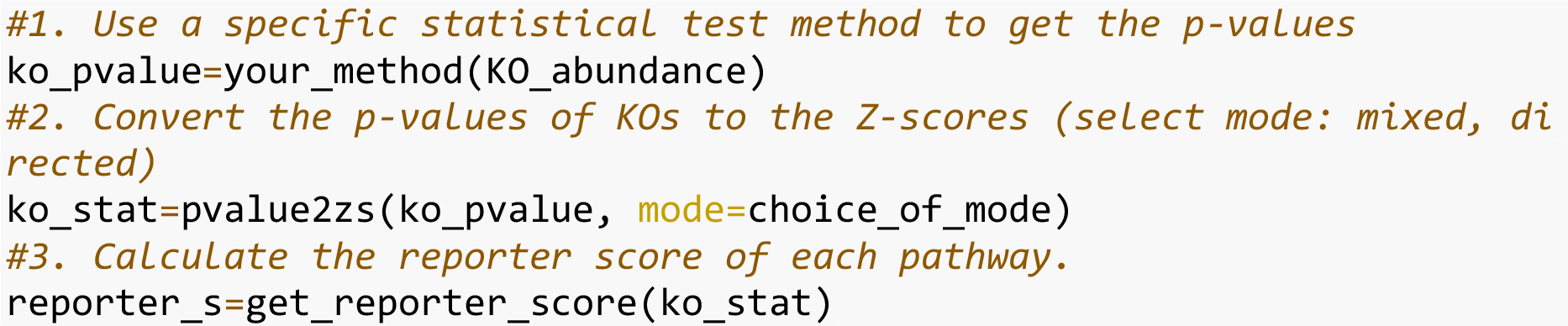

### Benchmark datasets

Benchmark datasets included one metagenomic KO profile (ex_KO_profile downloaded from https://github.com/wangpeng407/ReporterScore), 24 gene expression profiles of multiple human tissue types with disease, and 9 gene expression profiles of wild-type/knockout mice from the GEO database (https://www.ncbi.nlm.nih.gov/geo/).

We used these benchmark datasets with two-group or multi-group experimental designs to investigate the performance of GRSA, including similarities and differences between the two modes, statistical methods, and comparing GRSA with other commonly used enrichment analysis methods. Details of the datasets can be found in Supplementary Table 2 and Supplementary Table 3.

### Case study datasets

Three case studies were re-analyzed using ReporterScore to demonstrate the versatile applications of GRSA, including microbiome, transcriptome, and metabolome.

Skin microbiome data were generated by Wang et al. (2021)^23^. Using the shotgun method, they sequenced 822 skin samples and constructed the complete Human Skin Microbiome Gene Catalog (iHSMGC). A full KO profile based on the KEGG database was provided on the website (https://ftp.cngb.org/pub/SciRAID/Microbiome/humanSkin_10.9M/AbundanceProfile/IHSMGC.KO.normalization.ProfileTable.gz). Metadata with details about including body site, sex, age and cutotype were obtained via https://static-content.springer.com/esm/art%3A10.1186%2Fs40168-020-00995-7/MediaObjects/40168_2020_995_MOESM2_ESM.xlsx.

Transcriptomic data were extracted from the study by Liu et al. (2017)^30^. They investigated time-course transcriptomic profiling of cardiomyocyte differentiation derived from human hESCs and hiPSCs. The gene expression matrix is available at https://www.ncbi.nlm.nih.gov/geo/query/acc.cgi?acc=GSE85331.

Metabolomic data were generated by Liang et al. (2020)^31^. They analyzed the untargeted mass-spectrum data of 784 samples from 30 pregnant women and built a metabolic clock with five metabolites that time gestational age. The 264 identified level-1 and level-2 metabolites with HMDB IDs and their log2(intensity) can be found at https://ars.els-cdn.com/content/image/1-s2.0-S009286742030564X-mmc2.xlsx.

## Statistical analyses

All statistical analyses were done on the R 4.2.2 platform. The developed ReporterScore package (https://github.com/Asa12138/ReporterScore) was used for GRSA and visualization. The venn diagram and venn network diagram were drawn by the pcutils package (https://github.com/Asa12138/pcutils/).

To compare the performance of different statistical methods in the two-group experimental design, we defined a Jaccard similarity index:

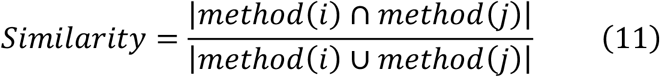

where method(i) (method(j)) is the number of significant pathways based on benchmark data sets enriched by different methods.

We used fuzzy c-means (FCM) clustering to explore the performance of different statistical test methods in multi-group experimental design (Extended Data Fig. b,c) and gene expression patterns in transcriptome (Fig. 4a). FCM is an unsupervised machine-learning technique that partitions a population into groups or clusters^42^. Three methods (Elbow, Silhouette, and Gap statistic) were used to determine the optimal number of clusters. In FCM, the membership score is the probability that a feature belongs to a cluster. Each feature is assigned to a cluster based on its highest membership score.

We compared GRSA against other most commonly used enrichment analysis methods. Fisher’s exact test is one of the most common functional enrichment analyses, which relies on an arbitrary cutoff of fold change and significance. GSEA is a classic functional class scoring method and analyzes all features based on their differential expression rank without prior feature filtering. GSEA calculates an Enrichment Score (ES) by moving through the ranked features list, increasing the ES if a feature is in the pathway, and decreasing the ES if not. These running sum values are weighted to amplify enrichment in the top- and bottom-ranking features. The ES is normalized to pathway size, yielding a Normalized Enrichment Score (NES). Positive and negative NES indicate enrichment at the top and bottom of the feature list, respectively. Lastly, a permutation-based *p-value* is computed, and multi-test correction is applied, yielding a False Discovery Rate (FDR) or Q value from 0 (significant) to 1 (not significant)^8^. However, the GSEA cannot be directly applied to multi-group or longitudinal datasets.

In the comparison of GRSA with other commonly used enrichment methods, we first calculated the *p-values* of features by t-test and performed adjustment of the *p-values* using the Benjamini & Hochberg (BH) method to control for False Discovery Rate (FDR). In order to perform the overrepresentation analysis, we selected the genes that have *p-values* less than 0.05. And the top 400 genes with the highest unsigned log-fold changes were considered as differentially expressed genes (DEGs)^43^. `fisher.test` (Fisher) was performed by the R base package. Improved `fisher.test` (CP) was performed by `enricher` function in the `clusterProfiler` package with the DEGs list. Gene set enrichment analysis (GSEA) was performed by the `GSEA` function in the `clusterProfiler` package, and the t-test statistic was used as the metric for ranking^44^ of `GSEA`. GRSA was performed with the BH-adjusted *p-values* of features by t-test. For convenience, the ReporterScore package provides an interface to the above-mentioned enrichment methods: `KO_fisher` for `fisher.test`, `KO_enrich` modified from `clusterProfiler` based on `fisher.test`, and `KO_gsea` modified from GSEA in `clusterProfiler`. These enrichment methods also support custom databases and are compatible with the format of the input data for the reporter_score function in GRSA, making it easy to make cross-comparisons.

Each human gene expression dataset in Supplementary Table 2 was associated with a disease with defined mechanism in a KEGG pathway (termed the target pathway). Ideally, a good enrichment analysis method would rank the target pathway at or near the top of the list with a small adjusted *p-value*^16^. We compared the performance of four enrichment analysis methods on these datasets.

For wild-type/knockout mice gene expression datasets (Supplementary Table 3), we considered the pathways containing the knockout gene as true positives and the pathways without the knockout gene as true negatives^16^. An adjusted *p-value* threshold of 0.05 was used to determine if a pathway is significantly affected. Based on the definitions of true positives (TP), true negatives (TN), false positives (FP), and false negatives (FN), we can calculate the sensitivity and specificity as follows:

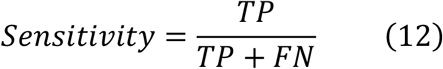

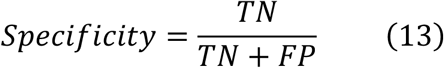

## Data availability

The source data for all figures are available from GitHub (https://github.com/Asa12138/GRSA_figures).

## Code availability

Code is available as an open-source R package ‘ReporterScore’, which can be downloaded from GitHub (https://github.com/Asa12138/ReporterScore). The main analysis scripts (Rmarkdown format) and source data are available from GitHub (https://github.com/Asa12138/GRSA_figures).

## Acknowledgments

We are grateful to our colleagues at the core facility of the Life Sciences Institute, especially the NECHO high-performance computing cluster. This research was partly supported by the grant from NSFC (82173645).

## Author contributions

C.P. and C.J. conceived the study. C.P. developed the R package. C.J. and C.P. collected all datasets. C.P. completed the main benchmarking and case study analyses. Q.C., S.T., and X.S. contributed to the analyses. C.P. and C.J. drafted and revised the manuscript with input from other authors.

## Additional information

**Supplementary Fig. 1-4**

**Supplementary Table 1-3**

