## Supplementary Materials for "Generalized Reporter Score-based Enrichment Analysis for Omics Data"

### Supplementary Figures

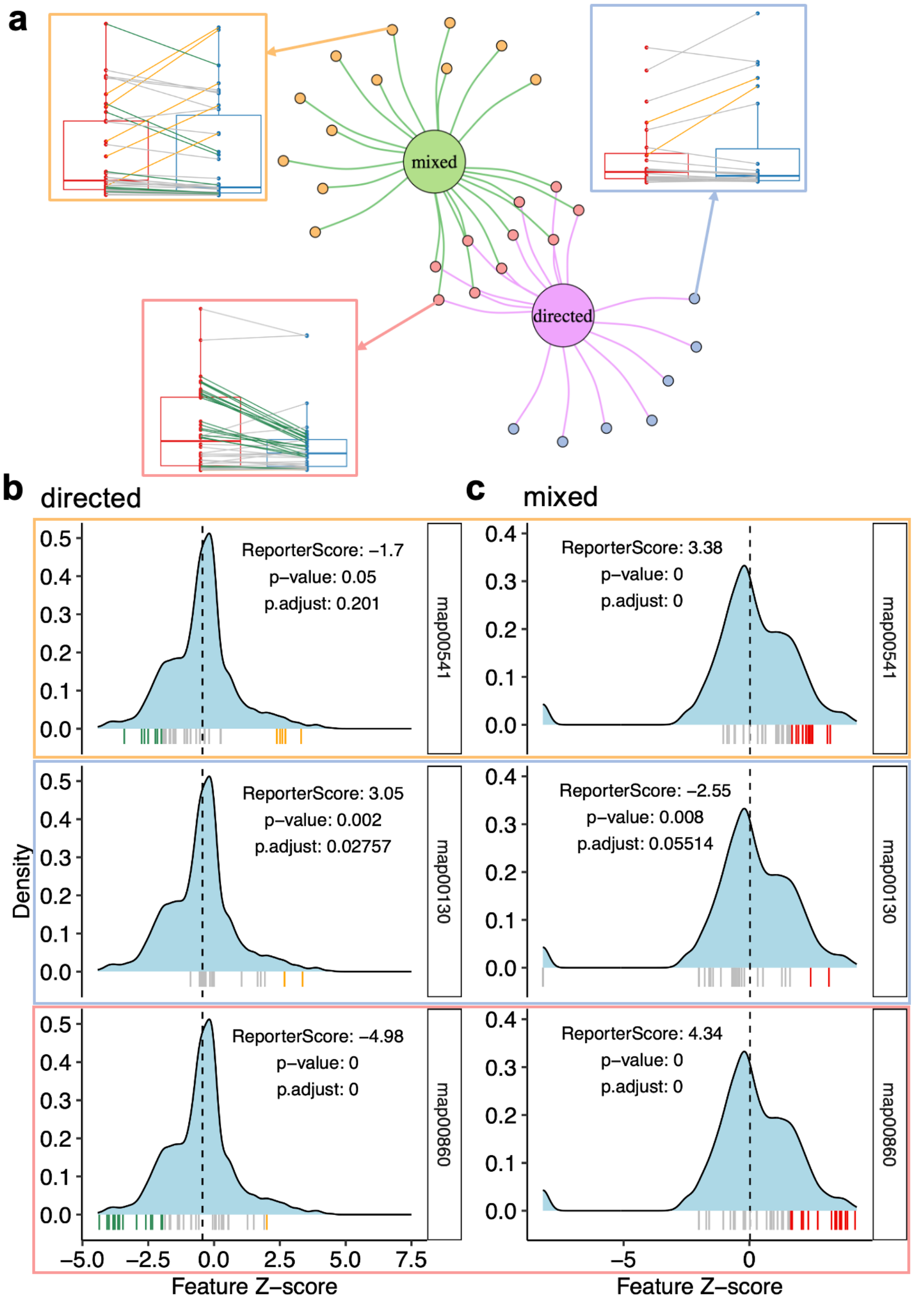

**Supplementary Fig. 1| Differences between the directed and mixed modes of GRSA results of the public ex_KO_profile dataset. a** Venn network of enrichment results using 'directed' and 'mixed' modes on the ex_KO_profile dataset. Each dot represents a significantly enriched pathway. The box charts show the trends of all KOs within selected pathways. Each line represents the trend of the average abundance of one KO. Line color indicates whether the KO is significantly enriched (orange), depleted (green), or neither (grey). **b**-**c** The distribution of KO Z-scores within the selected pathway compared to the background in the directed mode (**b**) and the mixed mode (**c**). The blue shading shows the density distribution of the background Z-score. The bottom whiskers represent the Z-scores of KOs within the selected pathway. The whisker color indicates whether the KO is significantly enriched (orange), depleted (green), or neither (grey).

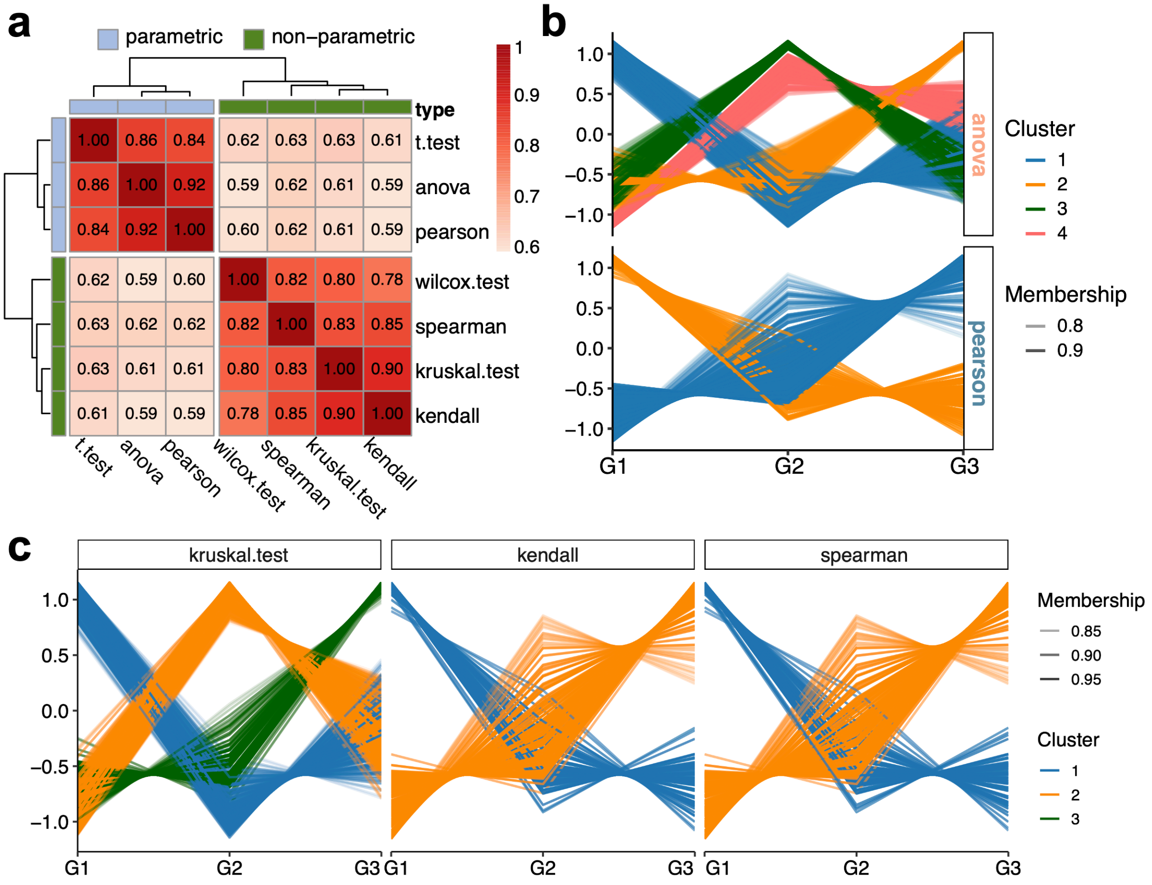


**Supplementary Fig. 2| Impact of the choice of statistical methods on the GRSA results.** **a** The heatmap shows the average similarity between enrichment results of parametric and non-parametric statistical methods on six benchmark datasets. The similarity was defined as the ratio of the number of pathways in the intersection to the union between the two methods. **b** C-means clustering of KO features, significance determined by the 'ANOVA' and 'Pearson' methods (parametric). **c** C-means clustering of KO features, significance determined by the 'Kruskal-Wallis test', 'Kendall', and 'Spearman' methods (non-parametric). The alpha (transparency) of each line reflects the value of its membership score, and the abundance was standardized. Only KOs within the significantly enriched pathways were used for the clustering. And only KOs with membership>0.8 were shown.

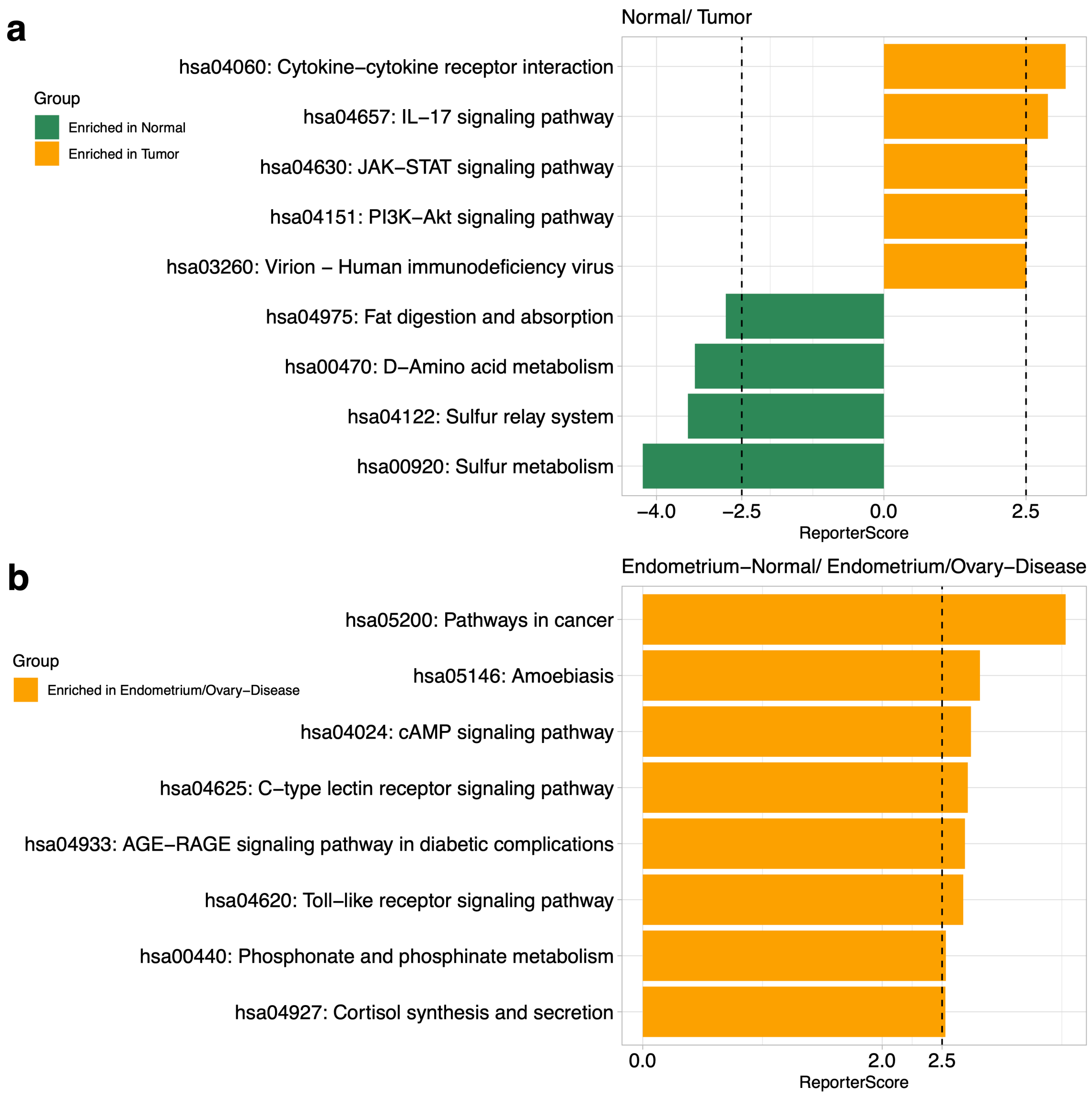


**Supplementary Fig. 3| GRSA-specific enriched pathways compared to GSEA.** The Bar chart shows GRSA-specific significantly enriched pathways compared to GSEA in GSE6344 (**a**) and GSE7305 (**b**); the threshold of 2.5 corresponds to a confidence of 0.995. Colors denote the enriched group.

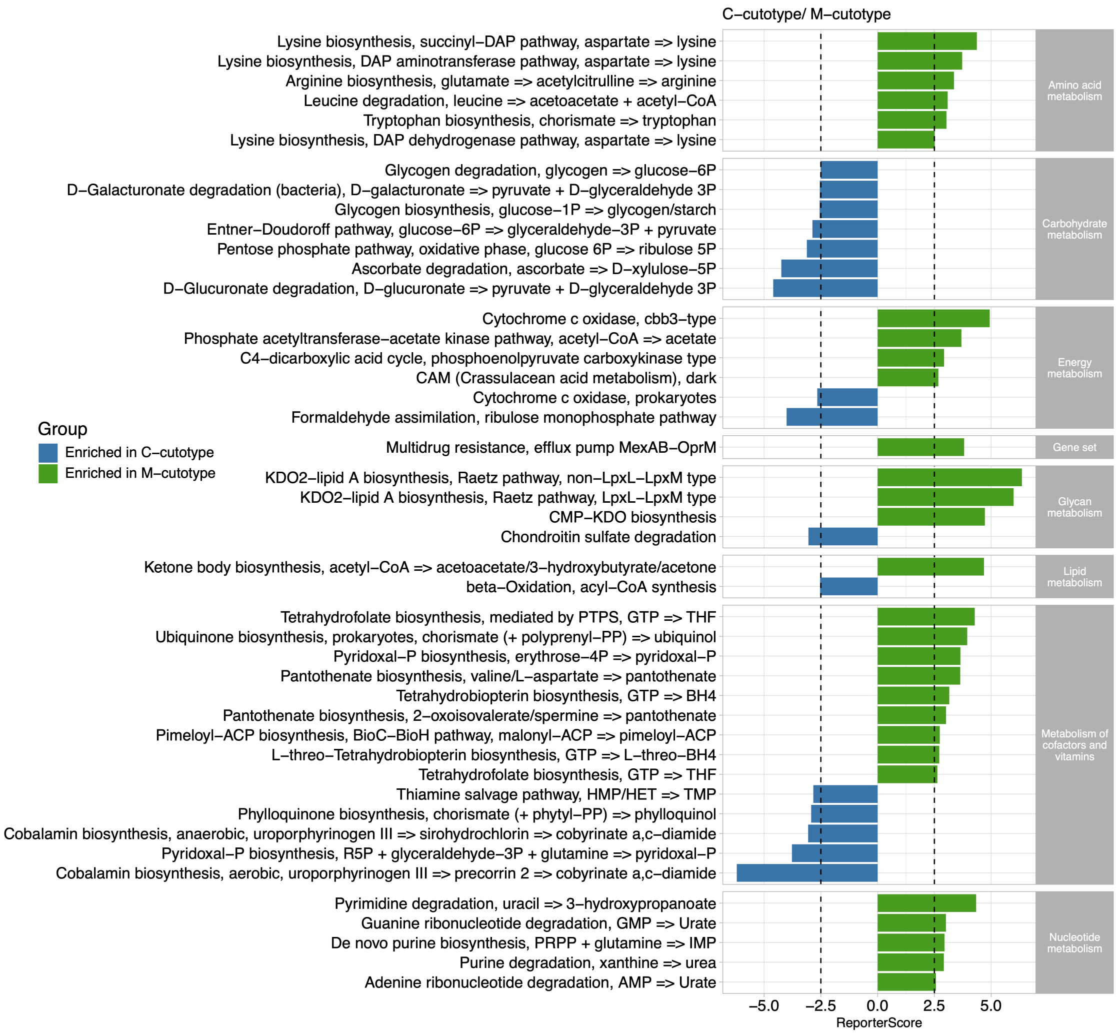


**Supplementary Fig. 4| Significantly enriched modules between *C-cutotype* and *M-cutotype*.** The threshold of 2.5 corresponds to a confidence of about 0.995, and these modules are grouped based on the KEGG level B.

### Supplementary Tables

**Supplementary Table 1**. Statistical methods supported by GRSA in the ReporterScore package.

| Method | Analysis | Type | Function |
|---|---|---|---|
| t-test | differential abundance | parametric | t.test() |
| Wilcoxon rank-sum test | differential abundance | non-parametric | wilcox.test() |
| ANOVA | differential abundance | parametric | anova() |
| Kruskal-Wallis rank-sum test | differential abundance | non-parametric | kruskal.test() |
| Pearson | correlation | parametric | cor.test(method='pearson') |
| Spearman | correlation | non-parametric | cor.test(method='kendall') |
| Kendall | correlation | non-parametric | cor.test(method='spearman') |

**Supplementary Table 2**. 24 benchmark gene expression datasets of 11 diseases used to compare enrichment analysis methods in this study.

| GEO id | Disease | Target pathway | Normal | Case | Pubmed | Tissue |
|---|---|---|---|---|---|---|
| GSE14762 | Renal cell carcinoma | hsa05211 | 12 | 9 | 19252501 | Kidney |
| GSE6344 | Renal cell carcinoma | hsa05211 | 10 | 10 | 17699851 | Clear cell RCC |
| GSE1297 | Alzheimer's disease | hsa05010 | 9 | 7 | 14769913 | Hippocampal CA1 |
| GSE5281EC | Alzheimer's disease | hsa05010 | 13 | 10 | 17077275 | Brain, Entorhinal Cortex |
| GSE5281HIP | Alzheimer's disease | hsa05010 | 13 | 10 | 17077275 | Brain, hippocampus |
| GSE5281VCX | Alzheimer's disease | hsa05010 | 12 | 19 | 17077275 | Brain, primary visual cortex |
| GSE65144 | Thyroid cancer | hsa05216 | 13 | 12 | 25675381 | Thyroid |
| GSE58545 | Thyroid cancer | hsa05216 | 18 | 27 | 26625260 | Thyroid |
| GSE41011 | Colorectal cancer | hsa05210 | 12 | 19 | NA | Colon |
| GSE55945 | Prostate cancer | hsa05215 | 7 | 12 | 19737960 | Prostate |
| GSE26910 | Prostate cancer | hsa05215 | 6 | 6 | 21611158 | Prostate |
| GSE8762 | Huntington's disease | hsa05016 | 10 | 12 | 17724341 | Lymphocyte |

| | | | | | | |
|---|---|---|---|---|---|---|
| GSE24250 | Huntington’s disease | hsa05016 | 6 | 8 | 21969577 | Venous cellular whole blood |
| GSE73655 | Huntington’s disease | hsa05016 | 7 | 13 | 26756592 | Subcutaneous adipose |
| GSE37517 | Huntington’s disease | hsa05016 | 5 | 8 | 22748968 | Neural stem cell |
| GSE14924 | Acute Myeloid Leukemia | hsa05221 | 10 | 10 | 19710498 | CD4 T Cell |
| GSE15471 | Pancreatic cancer | hsa05212 | 35 | 35 | 19260470 | Pancreas |
| GSE16515 | Pancreatic cancer | hsa05212 | 15 | 15 | 19732725 | Pancreas |
| GSE28735 | Pancreatic cancer | hsa05212 | 45 | 45 | 23918603 | Pancreas |
| GSE20153 | Parkinson’s disease | hsa05012 | 8 | 8 | 20926834 | Blymphocytes from peripheral blood |
| GSE19587 | Parkinson’s disease | hsa05012 | 10 | 12 | 20837543 | Brain |
| GSE26887 | Type II diabetes mellitus | hsa04930 | 5 | 7 | 22427379 | Left ventricle |
| GSE7305 | Endometrial cancer | hsa05213 | 10 | 10 | 17640886 | Endometrium/Ovarian tissue |
| GSE36389 | Endometrial cancer | hsa05213 | 7 | 13 | NA | Endometrium |

**Supplementary Table 3**. Nine knockout benchmark gene expression datasets used to compare methods in this study.

| GEO id | Knockout gene | Impacted pathway number | Normal | Case | Pubmed | Tissue |
|---|---|---|---|---|---|---|
| GSE22873 | Myd88 | 24 | 11 | 8 | 22075646 | Liver |
| GSE70302a | Il1a | 21 | 4 | 4 | 26224856 | Spinal cord |
| GSE70302b | Il1b | 41 | 4 | 4 | 26224856 | Spinal cord |
| GSE58120 | Il2 | 20 | 6 | 6 | 25652593 | Myeloid dendritic cells |
| GSE46211 | Tgfbr2 | 23 | 12 | 6 | 24496627 | Anterior palatal tissue |
| GSE138957 | Akt1 | 97 | 3 | 3 | 32937140 | Cell lines |
| GSE85754 | Fgfr1 | 16 | 4 | 4 | 28433771 | Breast |
| GSE88799 | Crebbp | 28 | 4 | 5 | 28069569 | B cell |
| GSE4451 | Met | 21 | 6 | 6 | 16710476 | Liver |
